# The gonad as a mediator of life history tradeoffs: Antagonistic hormonal pleiotropy facilitates evolutionary divergence in reproductive strategies

**DOI:** 10.64898/2026.07.06.736893

**Authors:** Victoria S. Farrar, Salvi Patel, Alexandra Sumarli, Kieran Samuk, Alison M. Bell

**Affiliations:** Department of Evolution, Ecology and Behavior, School of Integrative Biology, University of Illinois Urbana-Champaign, Urbana, IL, USA; Department of Evolution, Ecology, and Organismal Biology, University of California, Riverside, Riverside, California, USA; Carl R. Woese Institute for Genomic Biology, University of Illinois Urbana Champaign, Urbana, IL, USA

**Author notes:** Corresponding author. 505 S. Goodwin Ave., Department of Evolution, Ecology and Behavior, University of Illinois at Urbana-Champaign, Urbana, IL, 61801.

**Keywords:** RNA-Seq, spermatogenesis, transcriptomics, gonads, androgens, parental care, stickleback

## Abstract

High investment in current reproduction can limit future reproductive opportunities, but how selection shapes these hormone-mediated traits remains poorly understood. Androgens can mediate male reproductive investment, and in three-spined stickleback (*Gasterosteus aculeatus*), exert antagonistic effects on breeding effort versus spermatogenesis. To understand how shifts in reproductive strategy shape this tradeoff, we compared testes transcriptomes and androgen production between two recently diverged stickleback ecotypes that differ in reproductive strategy: the ancestral “common” ecotype, which provides paternal care, and the non-parental “white” ecotype, which has lost paternal care and prioritizes mating effort. During typical breeding, testes gene expression differed little between ecotypes. However, under prolonged summer-like conditions, testes gene expression diverged substantially. Common-biased genes were enriched for meiotic functions and spermatogenic cell type markers, suggesting commons had initiated spermatogenesis while whites had not. Instead, whites expressed higher levels of steroidogenic candidate genes and released significantly more 11-ketotestosterone than commons, indicating sustained investment in current reproduction. F1 hybrids released 11-ketosterone at intermediate rates, suggesting a genetic basis for this divergence. Sustained androgen production in whites may possibly delay the transition into spermatogenesis, limiting investment in future reproduction. These results illustrate how selection on hormonally-integrated traits can drive rapid divergence in life history strategy.

## INTRODUCTION

Life history theory predicts that greater investment in current reproduction limits future reproductive opportunities [1,2]. Animals experience reproductive tradeoffs both within seasons, such as the balance between parental care and mating effort during breeding [3], or across seasons, with overall reproductive effort incurring costs to growth and survival [4,5]. Individuals and populations can also vary in their intensity of reproduction across the lifespan, ranging from balanced, distributed reproductive investment across seasons to concentrated investment in the current reproductive bout [1,6]. In turn, intense reproductive investment can accelerate aging and shorten the lifespan, leading to a “faster” pace of life, and to an extreme, semelparity [4,7].

However, the proximate mechanisms shaped by selection on these reproductive strategies are not well understood. Hormones are potential mediators of life history tradeoffs due to their pleiotropic, and often antagonistic, effects on suites of traits across the lifespan [8–10]. Specifically, elevated androgens - which regulate male reproductive function and secondary sexual characteristics - can promote mating effort over parental care [11], accelerate senescence [12,13] and reduce long-term growth and survival [14,15]. Thus, high androgen levels during breeding may hamper investment in future reproduction.

How then does selection on androgen-mediated traits during breeding affect the traits androgens mediate at other life history stages? If androgen-mediated traits are integrated, theory predicts that selection on one reproductive trait could pull along correlated traits in other stages and hasten divergence in overall life history strategy [16,17]. Further, if antagonistic effects of androgens produce incompatibilities between discrete life history stages [8,11,18], high androgen levels may inhibit animals’ ability to transition to the next life history stage, leading to further divergence in the annual schedule. On the other hand, if traits in different life history stages can be regulated independently [16,19], the costs of elevated androgens may be circumvented from one stage to another [20]. For example, birds can maintain territorial aggression outside of the breeding season via neurosteroid conversion [21], thus avoiding the costs of high circulating androgens [22]. Parsing apart whether traits involved in reproductive investment are hormonally integrated or independent across life history stages is key to understanding how endocrine divergence can produce novel reproductive strategies, and comparisons of populations that differ in androgen-mediated traits can provide unique insights into this question [e.g., 23,24].

To this end, we capitalized on the reproductive diversity and well-characterized antagonistic role of androgens in the annual reproductive cycle of the three-spined stickleback fish (*Gasterosteus aculeatus;* hereafter, “stickleback”). Stickleback exhibit a seasonally dissociated breeding pattern, with gamete production and peak mating behavior separated in time [25,26]. As spring approaches and days lengthen and temperatures rise, male stickleback develop nuptial coloration and exhibit nest building, courtship and parental behavior. Toward the end of the breeding season, as days shorten and the temperature drops, males initiate spermatogenesis and store sperm over the winter until the next summer breeding season [27,28]. Males are therefore sperm limited during breeding and must allocate sperm across matings [29]. Stickleback vary in the age of first reproduction (one to two years), lifespan, and number of reproductive attempts within a breeding season [30]. For annual stickleback populations that die after their first breeding season (their second summer), spermatogenesis occurs after the first summer of life. For iteroparous populations, spermatogenesis occurs both after the first summer and after subsequent breeding seasons. As stickleback males cannot be in full breeding condition and undergo active spermatogenesis simultaneously [25], these states represent discrete life history stages.

Importantly, in stickleback, breeding and spermatogenesis are rendered incompatible via the antagonistic effects of androgens. 11-ketotestosterone (11-KT), the primary androgen in teleosts [31], is required for the development of male secondary sexual characteristics and reproductive behavior like courtship and nest fanning [32,33]. In post-breeding males, 11-KT inhibits spermatogenesis [33,34]. Consistent with the role of 11-KT in reproductive behavior, populations that vary in breeding behavioral repertoires - including parental care [35] - often also vary in 11-KT levels [24,36]. This population variation provides opportunities to probe how selection on androgen-mediated traits during breeding shapes other androgen-dependent functions across the life cycle.

Here, we examined how divergence in androgen-mediated reproductive traits may affect life history transitions in two marine stickleback ecotypes that have recently diverged in reproductive strategies. The Nova Scotian “white” stickleback, named for its conspicuous male breeding coloration, prioritizes mating effort over parental care. Relative to males of the widespread “common” ecotype, which is ancestral, male whites show intense zig-zag courtship displays towards females [37,38], at rates much higher than other stickleback populations [e.g., 39,40]. Whites can also maintain courtship responses to females for extended periods of time (up to 42 days, versus the typical four to five days parental males remain responsive to females after fertilization) [37]. White stickleback have also lost species-typical male care. Instead of fanning and attending embryos in their nest after mating, male whites immediately disperse the embryos out of their nest post-fertilization and return to vigorous courtship [37,38]. Male whites can therefore return to the mating pool sooner and mate more frequently than male commons, who do not mate while they are providing paternal care (seven to fifteen days) [41]. Increased mating opportunities for male whites ostensibly leads to greater intersexual competition and sexual selection in whites than in commons, potentially affecting sperm counts. Further, these ecotypes differ in androgen profiles; male whites maintain higher whole body 11-KT concentrations post-fertilization, unlike male commons which show a decline 11-KT during peak parental care [36]. Despite these behavioral and physiological differences, white and common sticklebacks are very closely related and capable of producing fertile hybrids, yet remain genetically distinct [38,42–44]. This very recent divergence (i.e., within the last 10,000 years [42,45]) allows us to identify ecotype differences related to reproductive strategy rather than confounding species differences accumulated over evolutionary time.

In this study, we investigated 1) whether the high investment in mating effort in white stickleback is mediated by testes androgen production, 2) for how long androgen-mediated investment in current reproduction in whites can be maintained under prolonged exposure to cues that indicate favorable breeding conditions, and 3) whether whites’ high investment in current reproduction compromises investment in future reproduction, such as the transition into spermatogenesis. To gain insight into both androgen and sperm production, we analyzed testes transcriptomes in both ecotypes when they were either actively breeding or held on prolonged summer breeding conditions (long photoperiod and high temperature). Given that: 1) androgens antagonistically influence reproductive behavior and post-breeding spermatogenesis in male sticklebacks more generally [33,34], and 2) male whites and commons differ in androgen profiles [36], we predicted whites would show elevated capacity for steroid production (i.e., higher steroidogenic enzyme gene expression in testes) than commons during breeding. Additionally, since whites presumably mate more frequently and experience more mate competition than commons, we predicted testes gene expression would reflect higher sperm counts in breeding whites. If whites maintain elevated capacity for androgen production under prolonged summer conditions, we hypothesized this would preclude their ability to begin spermatogenesis. We also measured waterborne 11-KT release under prolonged summer conditions to test whether white and common ecotypes differ in androgen production when held on prolonged summer conditions. Hormones were measured in whites, commons, and F1 hybrids to functionally test predictions about androgen production capacity and to gain insights into the underlying genetic architecture. All studies were carried out in animals reared in a common laboratory environment to control for environmental effects.

## METHODS

### Experiment 1: Testes RNA sequencing Animals and tissue collection

Testes were collected from 18 male three-spined stickleback (*Gasterosteus aculeatus*) between June and November 2023. All males were reared in the laboratory but descended from wild-caught grandparents. Parental commons were descended from fish caught in Cherry Burton Road, New Brunswick, Canada (N 46° 01.516’, W 64° 06.150’), and non-parental whites descended from fish caught in Canal Lake, Nova Scotia, Canada (N 44° 29’54.0”, W 63° 54’09.1”). All fish were fed daily frozen bloodworms, mysis shrimp and copepods *ad libitum* and kept on 10 ppt salinity during experiments.

We sampled males of both white and common ecotypes during the summer breeding season (June through July) and post-breeding (October through November) (Figure 1A). Breeding males (n = 4 commons, 5 whites) were kept in summer photoperiod and temperature (16L:8D, 16.7) and had been on those conditions for an average of 23 (range:11-39) days upon tissue collection. Males with full red nuptial breeding coloration were housed in individual tanks (36L x 33W x 24H cm) and provided filamentous algae and sand as nesting materials. All breeding males were in a nesting state (as defined by [36,38]),as they had built nests and exhibited nest-oriented courtship behaviors when exposed to a gravid female ≥24 hours before sampling, but had never fertilized eggs.

**Figure 1.**
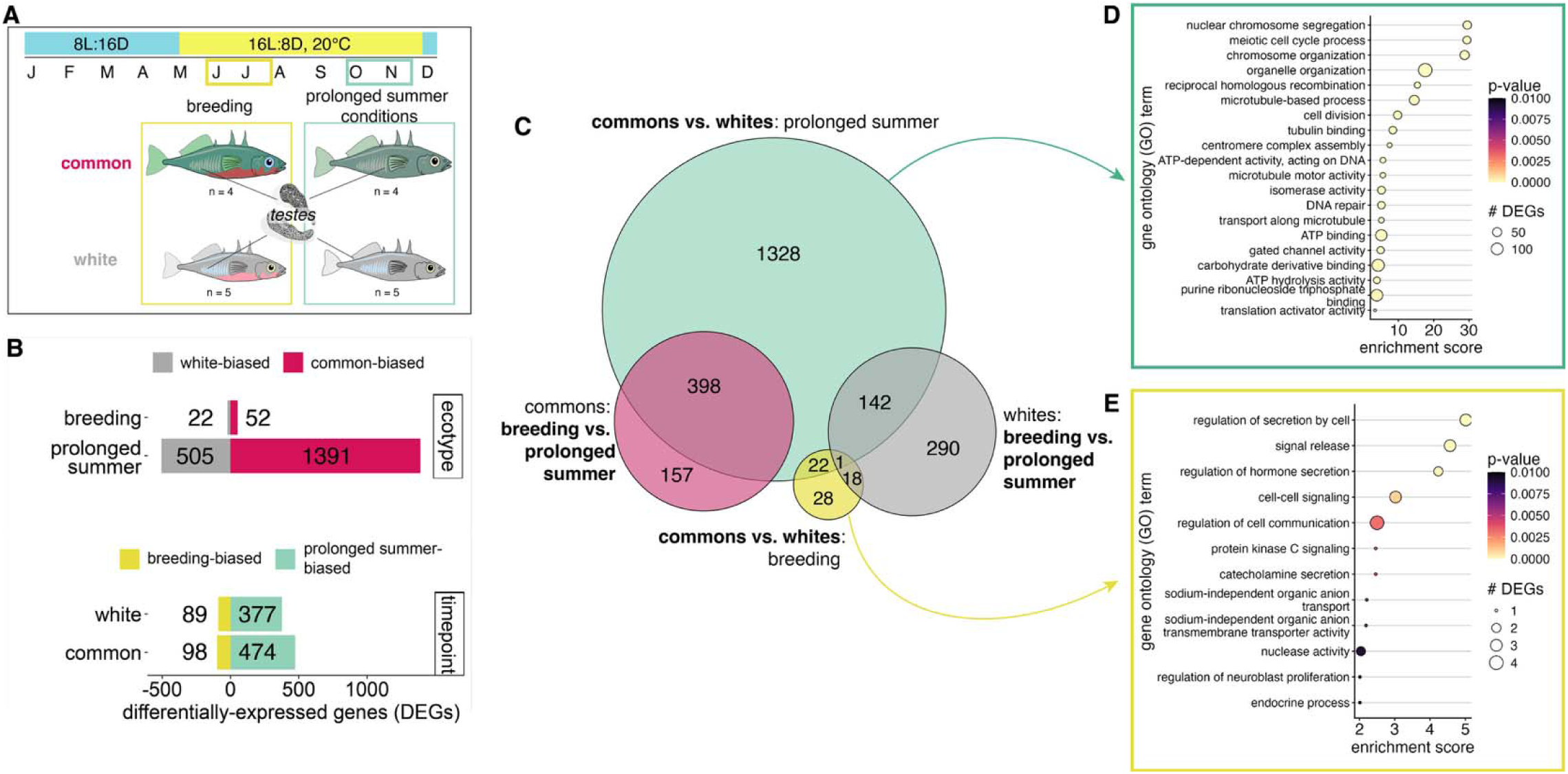
The testes of parental common and non-parental white ecotypes of stickleback are more transcriptionally different under prolonged summer conditions than during breeding. (**A**) Sampling design across laboratory photoperiod and temperature conditions over the year. (**B**) There are more differentially expressed genes (DEGs) between ecotypes under prolonged summer conditions versus during breeding (top). The ecotypes exhibit similar numbers of DEGs between sampling timepoints (bottom). (**C**) Euler plot showing overlap of DEG sets between pairwise contrasts. Note: 15 genes overlapping between timepoints in commons and in whites are not shown due to plot constraints. For all DEG set intersections, see UpSet plot (Supplemental Figure 2). More gene ontology (GO) terms were significantly enriched (Fisher’s exact test, *p* < 0.01) for (**D**) DEGs between ecotypes under prolonged summer as compared to (**E**) DEGs between ecotypes during breeding. Point size represents the number of DEGs annotated for each term, color represents significance.

Males held on prolonged summer conditions (16L:8D, 20) were sampled after they showed reduced signs of continued breeding in the lab; i.e., will no longer build nests when provided nesting material, and exhibit fading nuptial coloration. These males (n = 4 commons, 5 whites) were housed in communal tanks (53 L x 33 W x 24 H cm) with siblings. At the time of collection, males had been held on prolonged summer photoperiod and conditions (16L:8D, 20) for an average of 143 (range:109-167) days total. Previous studies have defined males as “post-breeding” when held on these conditions for 51-142 days [e.g., 27,46,47].

At both timepoints, fish were euthanized via rapid decapitation and both testes were dissected out, flash frozen and stored at −80 for further processing. All tissues were collected between 08:00-12:00. All procedures were approved by the XXX Institutional Animal Care and Use committee (IACUC) protocol #21031.

### RNA extraction and sequencing

To extract total RNA, both testes from each fish were homogenized together using ceramic beads (2.8mm, Fischer Scientific, cat #15-340-154), then extracted and purified using Purelink RNA Mini kits (Invitrogen, cat #12183018A). RNA was quantified using Qubit RNA Broad Range Assay (Invitrogen, cat #Q10210) and quality assessed using an Agilent 2100 Bioanalyzer. Libraries were prepared with Kapa Hyper Stranded mRNA library kit (Roche), then sequenced on two NovaSeq X Plus 25B lanes for 151 cycles from both ends of the fragments (paired end reads). Libraries were prepared and sequenced at the XXX center at XXX. Sequencing yielded an average of 58.5 million reads per sample per sequencing direction. Fastq files were demultiplexed using the *bcl2fastq* (v2.20, Illumina), then quality was assessed using *fastp* (v.0.23.4). We then aligned reads to the *Gasterosteus aculeatus* V5 reference genome (Ensembl release version 112) using *STAR* (v.2.7.10). Indexed and sorted reads were counted using featureCounts in *Subread* (v.2.0.4).

### Analysis

We then analyzed differential gene expression using *edgeR* v.4.4 [48] in R (v. 4.3.3). We filtered genes to include those with > 0.5 counts per million in four or more samples, which resulted in 21,118 total genes. Gene counts were normalized to trimmed mean of M-values. To identify differentially expressed genes, we conducted pairwise comparisons between ecotypes at each timepoint and between timepoints within ecotypes. We used a tagwise dispersion estimate after computing common and trended dispersions, then called differential expression using a “glm” approach. Genes with FDR < 0.05 were considered significantly differentially-expressed genes (DEGs). We visualized DEG set overlap using *eulerR* [49]. We analyzed gene ontology (GO) enrichment for all DEG sets using Fischer’s exact test in *topGO* [51] (enrichment called at *p* < 0.01) and reduced enriched GO annotation lists based on semantic similarity with *rrvgo* [52].

Given our hypothesis that androgen regulation has diverged between ecotypes, we focused on candidate genes involved in steroidogenesis or steroid hormone response (Table S1). As these candidates were defined *a priori*, we report *p* values for their differential expression between pairwise contrasts from edgeR with a Benjamini-Hochberg (FDR) correction applied specifically to the candidate gene set.

To examine if differences in testes gene expression might be due to differences in androgen receptor activity, we tested for enrichment of predicted androgen receptor target genes in our DEG lists. Teleosts have two androgen receptor genes: *ar1* and *ar2*, which code for proteins AR, and ARβ respectively [53]. To define predicted androgen receptor target genes, we built a gene regulatory network using GENIE3 v.1.33 [54]. GENIE3 uses a random forest approach to predict the expression of target genes based on the expression of genes defined as “regulators”, and has been shown to accurately predict transcription factor targets when functionally tested [55]. Importantly, however, GENIE3 does not directly predict receptor binding. We validated the GENIE3 predicted targets of androgen receptors by building a gene regulatory network for brain RNA-Seq data from sticklebacks treated with 11-ketoandrostenedione (11-KA), the precursor to the main teleost androgen 11-ketotestosterone [56], and found that GENIE3-predicted *ar2* target genes were significantly more likely to be enriched in 11-KA biased genes than other transcription factors (Supplemental Methods; Figure S1), consistent with the hypothesis that higher 11-KT leads to higher ARβ activation. Because our validation showed that predicted target genes of *ar2*, but not *ar1*, were enriched in genes upregulated by 11-KA treatment (Figure S1), and ARβ preferentially binds 11-KT in stickleback [57], we tested for enrichment of only predicted *ar2* target genes in our data.

To build a gene regulatory network from the testes gene expression data, we included all stickleback transcription factors from the Animal Transcription Factor database 4.0 [58] as regulators, as in Barbasch et al.[59].Predicted *ar2* target genes were defined by filtering the top 1,000,000 highest weighted regulatory links across all transcription factors (6.1% of all network links) down to those with *ar2* (ENSGACG00000020332) as the regulatory gene, yielding 1,403 predicted *ar2* target genes. We then tested for significant enrichment of these predicted *ar2* target genes across our DEG lists using Bonferroni-corrected hypergeometric overlap tests.

To further test whether stage of spermatogenesis differed between ecotypes on prolonged summer conditions, we examined whether our DEGs lists were enriched with 3,694 genes differentially expressed in specific testis cell clusters (e.g., spermatocytes or spermatogonia) in a single nuclei RNA-Seq dataset generated from male stickleback actively undergoing spermatogenesis [60]. We tested for enrichment of the differentially expressed marker genes unique to each of the 15 cell clusters identified in Shaw et al. [60] in our list of DEGs between ecotypes on prolonged summer conditions using Bonferroni-corrected hypergeometric overlap tests.

### Experiment 2: Waterborne hormone analysis

To test whether white and common ecotypes indeed differ in androgen production when held on prolonged summer conditions, we collected waterborne hormone samples from another set of males held on summer conditions for a similarly extended period. Additionally, to identify any potential genetic bases to androgen production, we also sampled F1 hybrids between whites and commons generated through reciprocal crosses.

### Animals and waterborne hormone collection

Waterborne hormone samples were collected in November 2024 from males (n = 56) held on prolonged summer conditions for an average of 162 (range:154 –175) days total. As in the previous experiment, males exhibited little to no nuptial coloration, were no longer territorial and were communally housed in large tanks (53 L x 33 W x 24 H cm) with siblings. All fish were fed daily frozen bloodworms, mysis shrimp and copepods *ad libitum*, and were held on summer photoperiod and temperature (16L:8D, 20) at 10 ppt salinity. All samples were collected over 21days.

Fish sampled were either commons (n = 17), whites (n = 9), or F1 hybrids with common fathers (WxC; n = 14) or white fathers (CxW; n =16). All fish in this experiment were generated as embryos via artificial fertilization from wild-caught parents in June 2023 and reared in the laboratory at Institution XXX then shipped to Institution YYY as adults in May 2024 before the breeding season (as described in Behrens et al. [61]). Common parents originated from Blue’s Cove, Nova Scotia, Canada (45°53’55.7"N 61°05’11.1"W), and white parents from Salmon River, Nova Scotia, Canada (45°21’9.57"N 61°28’22.20"W). Reciprocal F1 crosses were generated by crossing gametes from these populations.

To collect waterborne hormones, individual fish were placed in opaque glass beakers containing 50 mL of clean water. After 30 minutes, fish were removed and the water sample was filtered with qualitative filter paper (11.0 cm, Double Rings). These methods have been previously used to measure androgen excretion rates in stickleback [62,63], and waterborne androgens positively correlate with plasma concentrations in stickleback [63]. Fish were then rapidly euthanized via decapitation and males confirmed through the presence of testes. Water samples were stored at −20 until steroid extraction.

### Steroid extraction

Steroid hormones were extracted from water samples using solid-phase extraction with Sep-Pak C18 cartridges (Waters Corp., cat. #WAT020515). Cartridges were activated with 6 mL of 100% methanol, conditioned with 6 mL of reverse osmosis-treated water, then experimental water samples were run through cartridges using vacuum manifold. Steroids were eluted with 4 mL of methanol, then dried under nitrogen gas at 37. Extracts were stored at −20.

### Enzyme-linked immunoassay (ELISA)

We measured 11-ketotestosterone in our samples as this is the primary androgen in male fish, including breeding stickleback [31], and whole body 11-KT concentrations significantly differ between white and common ecotypes during breeding [36]. To quantify waterborne 11-KT concentrations, we used the DetectX^TM^ 11-Ketotestosterone Enzyme Immunoassay (Arbor Assays, cat.# K079). We resuspended all dried steroid extracts in 0.45 mL of assay buffer before plating. All samples and standards were run in duplicate. Plates were prepared following the manufacturer’s guidelines and read at 450 nm A pooled sample was included on each plate to determine inter-assay %CV. All samples were run across two plates. The mean intra-assay %CV was 7.82, and inter-assay %CV was 5.31.

We interpolated 11-KT concentrations from the standard curve using MyAssays.com online software (myassays.com). For samples with concentrations exceeding 2000 pg/mL, the upper limit of the standard curve, concentrations were conservatively truncated to 2000 pg/mL (n = 2).

### Statistical analysis

All statistical analysis was performed using R (version 4.4.3; R Core Team, 2025). We calculated 11-KT release rate as 11-KT sample concentration per body mass per time held (ng/g/hr). As our data did not meet assumptions of normality even after transformation (Shapiro-Wilk normality test, *p* < 0.001), we used non-parametric statistical tests. To test whether 11-KT release rates differed between ecotypes, we ran a one-way Kruskal-Wallis test, followed by a post-hoc Dunn test with Bonferroni correction for multiple comparisons.

## RESULTS

### Testes gene expression differed between ecotypes under prolonged summer conditions

Male whites and commons exhibit dramatically different reproductive strategies during the breeding season. Therefore, we predicted that they would differ in testes gene expression patterns. Consistent with this hypothesis, testes gene expression differed between whites and commons during active breeding (Figure 1A). However, testes gene expression differed to an even greater degree between whites and commons held under prolonged summer conditions. Over 25 times more genes were differentially expressed between ecotypes under prolonged summer conditions (n =1896) than during active breeding (n = 74) (Fig.1B). 22 genes were differentially expressed between ecotypes at both timepoints (Fig.1C), representing possible constitutive ecotype differences in the testes. Both ecotypes showed relatively similar numbers of differentially expressed genes (DEGs) between timepoints (n = 572 in commons, 466 in whites), with most of these genes expressed at higher levels under prolonged summer conditions (Fig.1B). However, most genes that shifted significantly across timepoints were unique to ecotype, with only 15 DEGs shared between ecotypes.

### Spermatogenesis-related functions and cell type markers were enriched in testes DEGs under prolonged summer conditions

The large number of DEGs between ecotypes under prolonged summer conditions was significantly enriched for gene ontology (GO) biological processes involved in nuclear chromosome segregation, meiotic cell cycle processes, chromosome organization, reciprocal homologous recombination, cell division, among others (Fig.1D, Table S4). Together, these terms suggest large ecotype differences in active cell division and meiotic state, implying differences in spermatogenic activity. In contrast, the DEGs between ecotypes at breeding were enriched for regulation of cell secretion, including of hormones and catecholamines (Fig. 1E; Table S5), possibly representing differences in hormone or neurotransmitter regulation.

Further analyses revealed that the meiotic and spermatogenesis-related processes enriched in the prolonged summer DEGs were specifically upregulated in commons (Fig. 2A; Table S6). As these differences in expression might reflect differences in overall spermatogenic cell state of the testes rather than changes in transcriptional regulation within cell types, we leveraged a published testes single-nuclei RNA-Seq dataset [60] and tested for enrichment of DEGs identified between cell types involved in active spermatogenesis in stickleback males.. DEGs between ecotypes under prolonged summer conditions were significantly enriched for DEGs from early spermatogonia, and mid- and late-spermatocytes (Bonferroni-corrected hypergeometric overlap tests, p.adj < 0.01; Fig. 2B). Specifically, common-biased genes at this timepoint were enriched for genes characteristic of meiotic cell types (spermatocytes; as defined by [60]), while white-biased genes were enriched only for genes from early spermatogonia, a pre-meiotic cell type, suggesting that commons had begun meiosis under prolonged summer conditions while whites had not. In contrast, DEGs between ecotypes at breeding were not significantly enriched for any cell cluster markers. This enrichment of meiotic cell cluster genes provides further evidence that when held under prolonged summer conditions, the ecotypes differ in the cellular composition of their testes and their progression into spermatogenesis, resulting in differences in testes gene expression.

**Figure 2.**
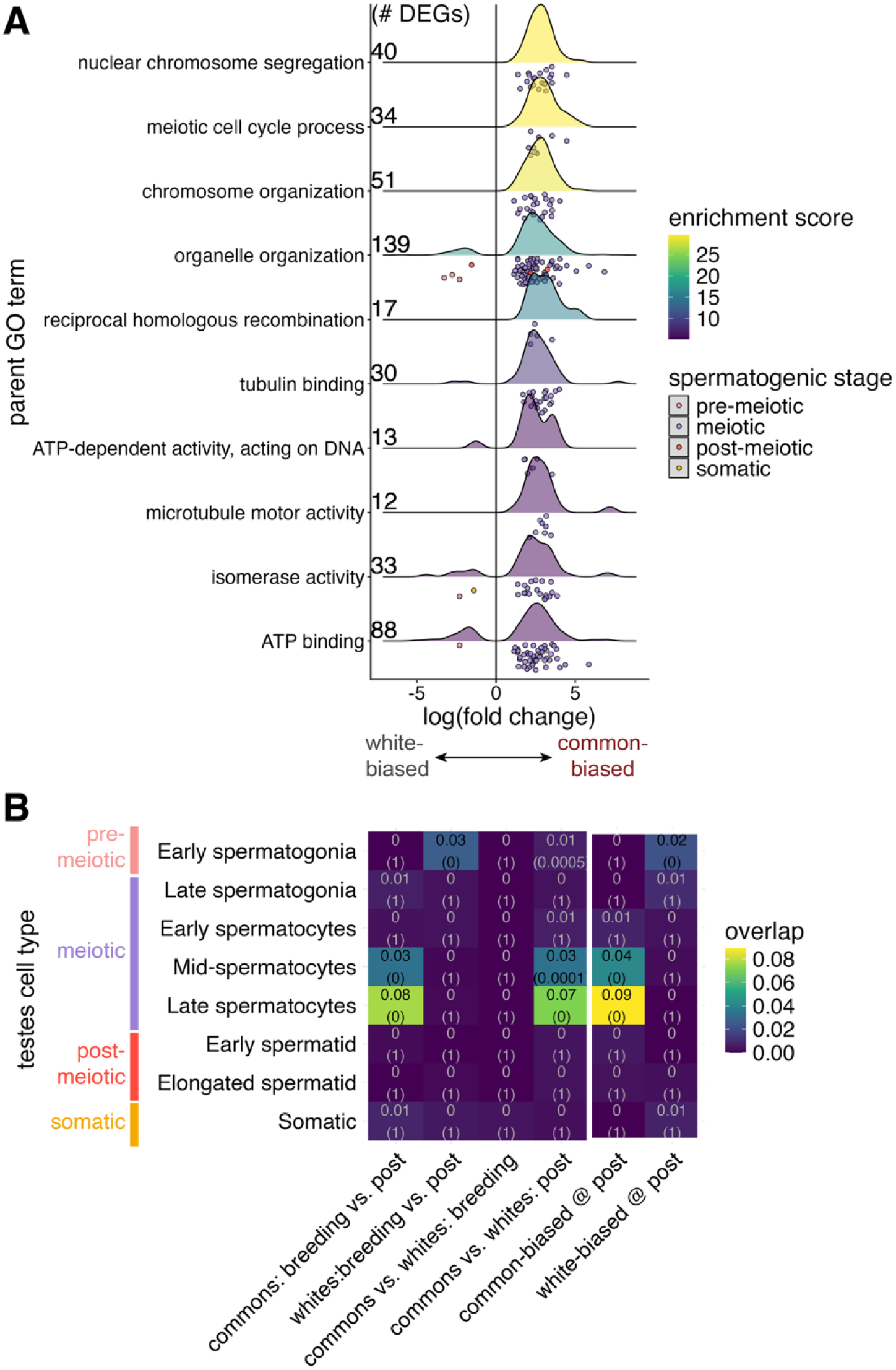
Common-biased DEGs in testes under prolonged summer conditions are enriched for spermatogenic cell types. (**A**) The top 10 most significantly enriched parent GO terms for DEGs between ecotypes under prolonged summer show log fold changes biased towards commons (positive log fold change). Density plot color represents GO term enrichment score. These GO terms include DEGs which are also differentially expressed between spermatogenic cell clusters in single nuclei RNA-Seq data from testis [60], shown as individual points (raincloud) under the density plots. Point color corresponds to the spermatogenic stage of the cell type cluster (pre-meiotic, meiotic, post-meiotic, or somatic) as assigned in [60]. (**B**) DEGs from meiotic cell types mid-spermatocytes and late-spermatocytes are significantly enriched in ecotype DEGs at prolonged summer and common-biased DEGs specifically (warmer colors indicate higher degree of overlap; intersection over union). Degree of overlap with Bonferroni-adjusted p-values from hypergeometric overlap tests are shown in parentheses. Common-biased DEGs and white-biased DEGs at prolonged summer are plotted separately to the right. Spermatogenic stage assignment and colors (left) are from [60].

### Steroidogenic candidate genes showed a greater decrease in expression in commons than whites under prolonged summer

The expression of multiple candidate genes along the steroidogenic pathway trended (0.05 < p.adj< 0.10) towards ecotypic differences. Of these, six were upregulated in whites compared to commons under prolonged summer conditions, including luteinizing hormone receptor (*lhcgr*), steroid acute regulator protein (*star*), and enzymes *cyp11a2*, *cyp17a1*, *hsd3b1*, and *cyp11c1* (Figure 3). Upregulation of these genes may allow whites to maintain relatively higher capacity for steroid production than commons under prolonged summer. No steroidogenic candidate genes significantly differed between ecotypes in breeding males.

**Figure 3.**
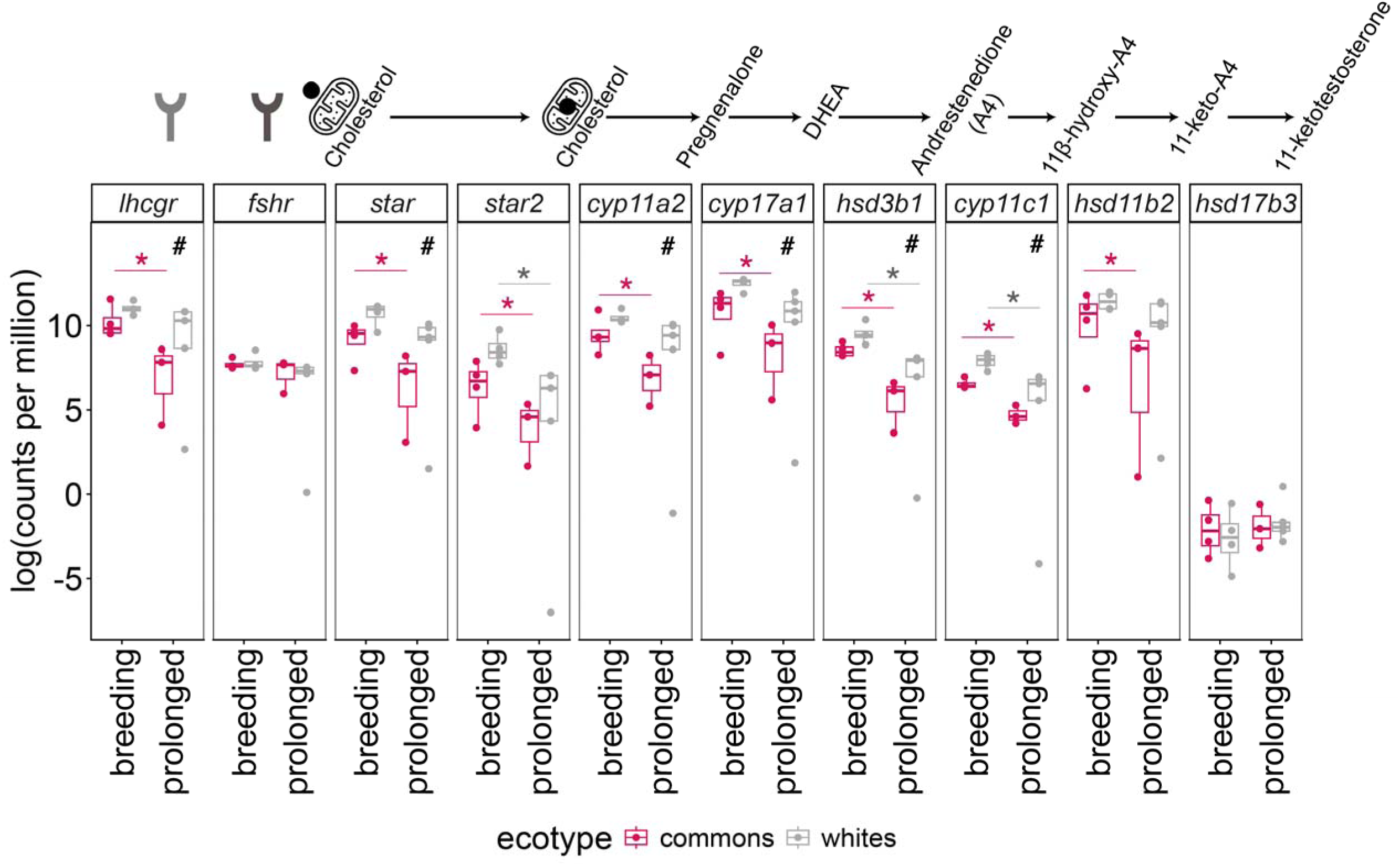
Candidate genes along the steroidogenic pathway are expressed at higher levels in whites compared to commons under prolonged summer conditions. Steroidogenic candidate genes differ between ecotypes on prolonged summer (“prolonged”) and between timepoints within ecotypes. Genes for gonadotropin receptors, steroidogenic cofactors and enzymes are plotted along the favored pathway of 11-ketotestosterone synthesis in teleosts [64,65], including stickleback [45]. First, gonadotropins – especially luteinizing hormone [66] – bind their receptors and activate steroidogenic acute regulatory (StAR) protein, facilitating the transport of cholesterol to the inner mitochondrial membrane for conversion by steroidogenic enzymes. Then, cytochrome p450 family 11 subfamily a (CY11A2) converts cholesterol to pregnenolone, which cytochrome p450 family 17 subfamily a (CYP17A1) converts to dehydroandrostenedione (DHEA), which 11-beta-hydroxysteroid dehydrogenase family 3 (HSD3B1) converts to androstenedione (A4). A4 is then converted to 11-beta-hydroxy-androstenedione (11β-OH-A4) by cytochrome p450 family 11 subfamily c (CYP11C), the teleost homolog of mammalian CYP11B1 [67]. 11β-OH-A4 is then converted to 11-ketoandrostenedione (11-KA, here 11-keto-A4) by hydroxysteroid 11-beta dehydrogenase 2 (HSD11B2). 11-KA is then the main androgen produced by the testes in stickleback, which is released into circulation where it is primarily converted to 11-ketotestosterone (11-KT) by hydroxysteroid 17-beta dehydrogenase (HSD17B) in blood [68]. Box plots show the median and first and third quartiles, and the whiskers represent 1.5 times the interquartile range, and points represent log counts per million for each individual male. Point color indicates ecotype. Asterisks (*) indicates p.adj < 0.05 and # indicates 0.05 < p.adj < 0.10 with bold black for ecotype differences and red and gray for timepoint differences within commons and whites, respectively.

Within ecotypes, more steroidogenic genes significantly decreased in expression over time (p.adj < 0.05) under summer conditions within commons (8 genes: *lhcgr, star, star2*, and enzymes*, cyp11a2, cyp17a1, hsd3b1, cyp11c1,* and *hsd11b2*) versus whites (3 genes: *star2* and enzymes *hsd3b1*, and *cyp11c1*) (Figure 3). Between timepoints, all steroidogenic candidate genes showed an average 1.63 log fold decrease in commons, but only an average 0.93 log fold decrease in whites, further evidence that whites experience less reduction in steroidogenic capacity under prolonged summer conditions than commons.

### Androgen receptor target genes are enriched in white-biased DEGs under prolonged summer

Given that whites showed relatively higher steroidogenic gene expression in testes under prolonged summer, suggesting they may produce more androgens, we predicted that these higher androgens might trigger more changes in gene expression downstream of the androgen receptor n whites. To test this, we examined whether predicted target genes of androgen receptor beta (*ar2*), which preferentially binds 11-KT in stickleback [57], were enriched in our DEG sets.

Predicted ARβ (*ar2*) target genes were significantly enriched in ecotype DEGs under prolonged summer (Bonferroni-adjusted hypergeometric overlap test, *p*. adj < 0.001, Table S2), as were both genes upregulated in whites and commons at that timepoint (p.adj < 0.001). Ecotype DEGs at breeding were not enriched for predicted *ar2* targets (Table S2).

### Whites and F1 hybrids maintain higher 11-KT release on prolonged summer conditions

To test if androgen production indeed differed between ecotypes under prolonged summer, as suggested by testes steroidogenic candidate gene expression, we measured waterborne 11-KT release by whites and commons in a separate experiment. To identify any possible genetic components to androgen release, we also compared 11-KT release in reciprocal F1 hybrid\ crosses between whites and commons. After an average of 162 days on summer photoperiod and temperature, the ecotypes significantly differed in 11-KT release rates (Figure 4; Kruskal-Wallis chi-squared = 22.657, df = 3, p < 0.001). Commons released 11-KT at an average of 0.13 (range: 0.08 – 0.24) ng/g/hr, which was less than whites (mean: 1.79, range: 0.12 – 5.22 ng/g/hr), F1(CxW) (mean: 0.47, range: 0.12 – 2.73 ng/g/hr), and F1(WxC) hybrids (mean: 0.83, range: 0.094 – 6.82 ng/g/hr). Commons had significantly lower release rates than whites (Dunn post-hoc test, Z = −3.558 p.adj = 0.002) and both F1(CxW) (Z = −4.25, p.adj < 0.001) and F1(WxC) hybrids (Z = −3.05, p.adj = 0.014). Whites and F1 hybrids did not significantly differ from each other (all p.adj = 1).

**Figure 4.**
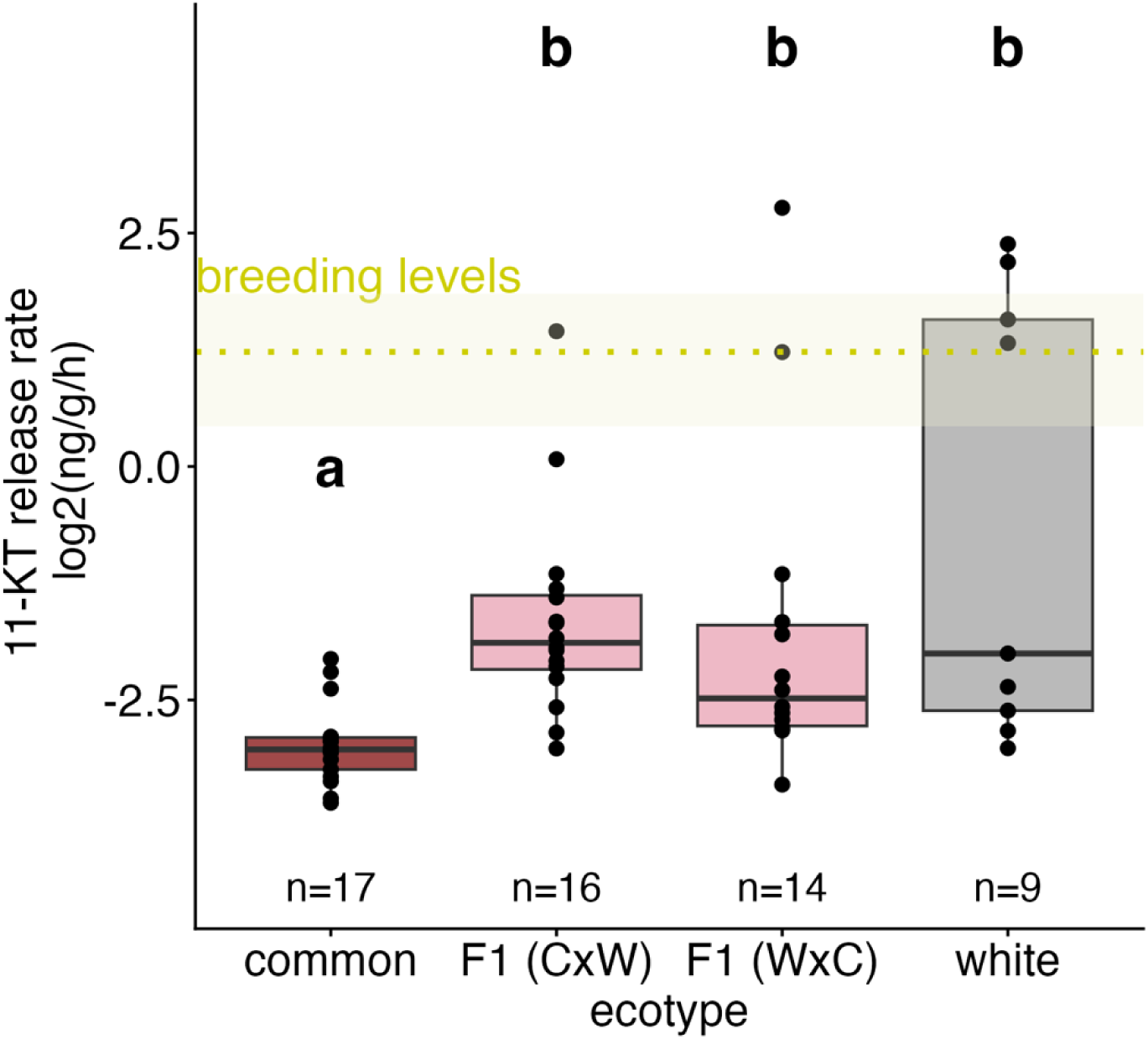
Male whites and F1 hybrids release more waterborne 11-ketotestosterone (11-KT) than commons under prolonged summer conditions. Log-transformed 11-KT release rates into water (ng/g body weight/hour) rank higher for whites and F1 hybrids than commons (Kruskal-Wallis chi-squared, *p* < 0.001). Box plots represent the median and first and third quartiles, with whiskers representing 1.5 times the interquartile range. Individual points represent values for individual males. Different letters represent significant pairwise differences between groups (Bonferroni-corrected post-hoc Dunn tests, *p* < 0.05). For reference, breeding levels of waterborne 11-KT from males in [63] are plotted, with the dotted yellow line representing the mean (2.23 ng/g/hr), and the yellow shaded region representing the full range (min = 1.35, max = 3.6 ng/g/hr) of 11-KT in breeding males in that study.

## DISCUSSION

Hormones can coordinate tradeoffs between current and future reproduction throughout the lifespan, but our understanding of how selection on life history strategies shapes endocrine regulation remains limited. Here, we leveraged two ecotypes of three-spined stickleback that have recently diverged in reproductive behavior and sex steroid profiles [36,38] to assess how this divergence has shaped androgen and sperm production in the testes, which are mutually-exclusive functions occurring during discrete life history stages in stickleback.

We predicted that during breeding, gonadal gene expression in the non-parental white ecotype would reflect increased steroid production capacity and higher sperm counts compared to the parental common ecotype. We also exposed males to prolonged summer conditions known to stimulate breeding (>100 days on long photoperiod, 16L:8D, and warm temperature, 20) to test for how long androgen-mediated investment in current reproduction can be sustained. We find that males of these ecotypes, when reared under shared laboratory conditions, show divergent testicular responses to prolonged summer conditions that are not apparent during breeding. Under prolonged summer conditions, genes from male commons were enriched for meiotic and recombination-related functions as well as spermatogenic cell type markers, indicating that commons’ testes had progressed into a spermatogenic cellular state, while whites’ testes had not. Whites’ testes also exhibited high levels of expression of steroidogenic candidate genes, suggesting that male whites may be able to sustain extended androgen production. To functionally test this, we measured waterborne 11-ketotesterone (11-KT) and found that whites, and F1 hybrids, released more 11-KT than commons under prolonged summer conditions. As androgens mediate incompatibilities between breeding behavior and gametogenesis in stickleback, our results highlight how heavy investment in current reproduction (mating effort) in white stickleback may shape their overall life history strategy.

Despite dramatic differences in behavior during the breeding season, we found relatively few differences in testes gene expression between ecotypes during breeding. Our results contrast with those from other stickleback species that exhibit divergent reproductive behavior, such as the Japan sea (*G. nipponicus*) and three-spined stickleback (*G.aculeatus*) species pair [24,39].

Japan sea males, which show more aggressive courtship behavior, also have higher plasma 11-KT and their testes express more LH receptors and steroidogenic enzymes during breeding [24]. However, despite striking differences in courtship behavior, whites and commons did not significantly differ in 11-KT concentrations at nesting [36]. This lack of difference in androgens at nesting is consistent with the lack of differences in testes steroidogenic candidate gene expression we observed. However, whites and commons do differ in 11-KT concentrations post-fertilization [36], with whites maintaining high levels while commons decline in 11-KT as they increase parental behavior. Comparing steroidogenic capacity in testes, as well as extra-gonadal 11-KT conversion in the blood [64], between the ecotypes *post*-fertilization may provide insight into within-breeding divergence in reproductive investment.

We hypothesized that selection may have favored increased sperm production in whites because they do not provide parental care and can thus return to the mating pool and mate more frequently than male commons. As stickleback have limited sperm stores during breeding, and the males in this study had never spawned, we specifically predicted that marker genes for spermatozoa (i.e., mature, motile sperm) would be enriched in DEGs between ecotypes at breeding. Instead, we found no sperm-related functions nor markers of any testis cell types were significantly enriched in breeding DEGs. However, mature spermatozoa are transcriptionally inactive [65], therefore our transcriptomic approach likely cannot directly detect differences in sperm counts. Further, spermatogenesis is quiescent during breeding (when 11-KT is high [27,28]), and the single nuclei RNAseq reference dataset was generated from fish undergoing active spermatogenesis [60] rather than from males with mature sperm. Therefore, the failure to detect enrichment for spermatogenic cell type markers in this study may not be surprising. A comparison of wild-caught males captured on the breeding grounds suggested that whites have smaller testes than commons even when correcting for body size [66]. This difference in testes size might reflect whites’ greater mating activity and release of a larger proportion of their limited sperm store rather than a lower initial investment in sperm. Ultimately, sperm counts and testis morphology will be required to address whether there are ecotypic differences in sperm.

In contrast to the results from the breeding timepoint, exposure to prolonged summer conditions produced large ecotypic differences in testicular correlates of spermatogenesis. Specifically, common-biased genes under these conditions were enriched for functions involved in meiosis and recombination, and for marker genes from mid- and late-spermatocyte clusters from the single nuclei RNASeq testes data [60]. Thus, differential testes gene expression between ecotypes during prolonged summer conditions may be attributable to differences in cellular composition or state. As high numbers of spermatocytes characterize active spermatogenesis in stickleback during the late summer/early fall, our data suggest commons had begun producing sperm under prolonged summer conditions, but whites had not. Instead, white-biased genes at this timepoint were enriched for markers of the pre-meiotic cell type early spermatogonia, which are present even when spermatogenesis is quiescent (i.e., breeding) [27,67]. Given that androgen treatment prevents male stickleback from initiating spermatogenesis post-breeding [33,34], the low signatures of spermatogenesis in whites may be partly due to sustained androgen levels under the prolonged summer conditions.

Indeed, we find multiple lines of evidence that whites maintain high androgen production capacity for a longer period than commons under prolonged summer conditions. First, male whites at this timepoint had higher testicular expression of genes known to correlate with higher plasma 11-KT in breeding stickleback, such as LH receptors, StAR, and steroidogenic enzymes [24], and few of these genes significantly decreased over time in whites. Second, white-biased genes at this timepoint were enriched for predicted targets of androgen receptor beta (*ar2*), the preferential receptor for 11-KT in stickleback [57], suggesting maintained androgen receptor activity. Third, whites released significantly more 11-KT than commons under prolonged summer conditions. Not only were white 11-KT release rates relatively higher on average, but there was a bimodal distribution within whites, with some individuals releasing11-KT at a rate on par with those of actively breeding stickleback in other studies (>1.5 ng/g/hr; [62,63]) while others released levels similar to commons. This variation in androgens within whites may have resulted from selection bias for fish with hints of breeding coloration under prolonged summer conditions or it may reflect true individual differences. Future studies with a larger sample size are needed to reveal how prevalent these high-androgen releasing individuals are within the white ecotype. Nonetheless, sustained androgen production would be consistent with whites’ broader strategy of high investment in current reproduction, including high levels of courtship behavior during breeding [37]. However, maintaining androgen production may hinder the onset of spermatogenesis and limit investment in future reproduction, as suggested by the low signatures of spermatogenesis we observed in the testes of male whites. Although whites had relatively higher 11-KT under prolonged summer conditions, we hesitate to interpret this as breeding readiness without validation through behavior or histology. The whites we sampled had little nuptial coloration and many released 11-KT at rates lower than breeding stickleback in other studies [63]. Rather than maintaining breeding, these whites may be delayed in their transition between life history stages, and correlates of spermatogenesis might have been detected if they had been sampled later.

We also find evidence that population differences in sustained androgen production have a genetic basis. Like whites, F1 hybrid crosses raised in the laboratory released higher 11-KT than commons under prolonged summer conditions, suggesting that the mechanisms maintaining androgen production may be partially dominant. That reciprocal hybrid crosses did not differ suggests no strong sex-linked effects on androgen maintenance, but undoubtedly many genetic components underlie steroid production and such genetic effects cannot be ruled out. During breeding, F1 hybrids show an intermediate phenotype between whites and commons in reproductive behaviors [38], which may reduce hybrid fitness and further isolate the ecotypes [61], but whether hybrids show similarly intermediate or dysregulated 11-KT release patterns during breeding is not known. Although we did not examine F1 hybrid testes, one might predict that hybrids’ testes would also show intermediate steroidogenic and spermatogenic gene expression under prolonged summer conditions. Future work linking genetic variation with differential gene expression in testes (i.e., expression quantitative trait loci) could uncover the genetic architecture underlying this divergence in androgen regulation.

One limitation to our approach is that we compare only a single source population of each ecotype for each experiment, thereby confounding potential population and ecotype differences. Recent population genomics studies reveal genetic structure within common populations across Nova Scotia [42], and whether these populations differ behaviorally or physiologically remains unknown. Importantly, however, we find a consistent pattern of relatively higher androgen production capacity in whites across two separate source populations and experiments, supporting a general ecotypic difference. Comparing breeding physiology across populations, including recently discovered stickleback populations in northern Scotland that convergently lost parental care (A. MacColl, *pers. comm.),* is a promising future direction for understanding the conserved mechanisms underlying shifts in reproductive strategies.

What proximate mechanisms might allow whites to continue androgen production and delay their transition into spermatogenesis under prolonged summer conditions? Reproductive opportunities for whites may be limited, partly because their high androgens may hasten reproductive senescence [12], and also because their conspicuous breeding coloration and behavior may increase predation risk (though this remains untested). Theory predicts that organisms with limited reproductive windows will modulate their reproductive effort less in response to environmental cues [18]. While the underlying neuroendocrine mechanisms are poorly understood, refractoriness in stickleback likely involves an endogenous timing mechanism that responds to photoperiod exposure [25,68]. Whites may have diverged in their sensitivity to these environmental cues and thus their thresholds for becoming refractory. Future experiments manipulating environmental conditions and comparing the responses of potential neural mediators, such as gonadotropin releasing hormone (GnRH) [47], between ecotypes could shed light on these mechanisms.

Over time, heavy investment in current reproduction (via mating effort) may lead whites to evolve a more annual, or semelparous, strategy. Stickleback can vary significantly in lifespan; most marine populations reproduce at two to three years of age and die after one to two breeding seasons, while others are annual, reproducing when one year old and dying after breeding [30]. While whites in the wild tend to be smaller than commons [43,66], which might suggest that they reproduce at one rather than two years of age, it remains unknown if whites are truly annual. Further, as we only studied males, we do not know if female whites also exhibit shifts in reproductive timing, paralleling shifts in other female white reproductive traits [69]. Demographic studies characterizing age at breeding and mortality in wild populations and in both sexes are required to confirm true ecotype differences in these life history traits.

Overall, our results illustrate how divergence in androgens, which exert antagonistic effects on two discrete life history stages - breeding and spermatogenesis - may contribute to rapid divergence in reproductive strategy between these two stickleback ecotypes. We observed these differences in life history stage transitions in the laboratory and in fish from multiple source populations and experimental years, pointing to a potential genetic basis for these ecotypic differences. Together, these results provide a novel example of how selection on hormonally-integrated traits can be a force for rapid adaptation and divergence in life history strategies.

## ACKNOWLEDGEMENTS

We thank Sarah Westrick and Anjali Tibudan for help validating steroid extractions, and Jun Kitano for sharing brain RNA-Seq data for GRN validation. Mike White and members of the Bell laboratory provided valuable comments.

## REFERENCES

1. Stearns SC. 1976 Life-history tactics: a review of the ideas. Quarterly Review of Biology 51, 3–47.

2. Roff DA. 1992 Costs of reproduction. In The evolution of life histories: theory and analysis, pp. 145–179. New York: Chapman & Hall.

3. Stiver KA, Alonzo SH. 2009 Parental and Mating Effort: Is There Necessarily a Trade-Off? Ethology 115, 1101–1126. (doi:10.1111/j.1439-0310.2009.01707.x)

4. Reznick D. 1985 Costs of Reproduction: An Evaluation of the Empirical Evidence. Oikos 44, 257. (doi:10.2307/3544698)

5. Harshman LG, Zera AJ. 2007 The cost of reproduction: the devil in the details. Trends in Ecology & Evolution 22, 80–86. (doi:10.1016/j.tree.2006.10.008)

6. Gaillard J-M, Pontier D, Allainé D, Lebreton JD, Trouvilliez J, Clobert J, Allaine D. 1989 An Analysis of Demographic Tactics in Birds and Mammals. Oikos 56, 59. (doi:10.2307/3566088)

7. Stearns SC. 1977 The Evolution of Life History Traits: A Critique of the Theory and a Review of the Data. Annu. Rev. Ecol. Syst. 8, 145–171. (doi:10.1146/annurev.es.08.110177.001045)

8. Ricklefs RE, Wikelski M. 2002 The physiology/life-history nexus. Trends in Ecology & Evolution 17, 462–468. (doi:10.1016/S0169-5347(02)02578-8)

9. Zera AJ, Harshman LG. 2001 The Physiology of Life History Trade-Offs in Animals. Annual Review of Ecology and Systematics 32, 95–126. (doi:10.1146/annurev.ecolsys.32.081501.114006)

10. Finch CE, Rose MR. 1995 Hormones and the Physiological Architecture of Life History Evolution. The Quarterly Review of Biology 70, 1–52. (doi:10.1086/418864)

11. Wingfield JC, Hegner RE, Dufty, AM, Ball GF. 1990 The ‘Challenge Hypothesis’: Theoretical Implications for Patterns of Testosterone Secretion, Mating Systems, and Breeding Strategies. The American Naturalist 136, 829–846. (doi:10.1086/285134)

12. Simons MJP, Sebire M, Verhulst S, Groothuis TGG. 2021 Androgen Elevation Accelerates Reproductive Senescence in Three-Spined Stickleback. Front. Cell Dev. Biol. 9, 752352. (doi:10.3389/fcell.2021.752352)

13. Heidinger BJ, Slowinski SP, Sirman AE, Kittilson J, Gerlach NM, Ketterson ED. 2022 Experimentally elevated testosterone shortens telomeres across years in a free-living songbird. Molecular Ecology 31, 6216–6223. (doi:10.1111/mec.15819)

14. Reed WL, Clark ME, Parker PG, Raouf SA, Arguedas N, Monk DS, Snajdr E, Nolan Jr. V, Ketterson ED. 2006 Physiological Effects on Demography: A Long-Term Experimental Study of Testosterone’s Effects on Fitness. The American Naturalist 167, 667–683. (doi:10.1086/503054)

15. John-Alder HB, Cox RM, Haenel GJ, Smith LC. 2009 Hormones, performance and fitness: Natural history and endocrine experiments on a lizard (Sceloporus undulatus). Integrative and Comparative Biology 49, 393–407. (doi:10.1093/icb/icp060)

16. Ketterson ED, Atwell JW, McGlothlin JW. 2009 Phenotypic integration and independence: Hormones, performance, and response to environmental change. Integrative and Comparative Biology 49, 365–379. (doi:10.1093/icb/icp057)

17. McGlothlin JW, Ketterson ED. 2008 Hormone-mediated suites as adaptations and evolutionary constraints. Phil. Trans. R. Soc. B 363, 1611–1620. (doi:10.1098/rstb.2007.0002)

18. Jacobs JD, Wingfield JC. 2000 Endocrine Control of Life-Cycle Stages: A Constraint on Response to the Environment? The Condor 102, 35–51. (doi:10.1093/condor/102.1.35)

19. Hau M. 2007 Regulation of male traits by testosterone: implications for the evolution of vertebrate life histories. BioEssays 29, 133–144. (doi:10.1002/bies.20524)

20. Wingfield JC, Lynn SE, Soma KK. 2001 Avoiding the ‘Costs’ of Testosterone: Ecological Bases of Hormone-Behavior Interactions. Brain Behav Evol 57, 239–251. (doi:10.1159/000047243)

21. Pradhan DS, Newman AEM, Wacker DW, Wingfield JC, Schlinger BA, Soma KK. 2010 Aggressive interactions rapidly increase androgen synthesis in the brain during the non-breeding season. Hormones and Behavior 57, 381–389. (doi:10.1016/j.yhbeh.2010.01.008)

22. Soma KK, Scotti M-AL, Newman AEM, Charlier TD, Demas GE. 2008 Novel mechanisms for neuroendocrine regulation of aggression. Frontiers in Neuroendocrinology 29, 476–489. (doi:10.1016/j.yfrne.2007.12.003)

23. Bergeon Burns CM, Rosvall KA, Hahn TP, Demas GE, Ketterson ED. 2014 Examining sources of variation in HPG axis function among individuals and populations of the dark-eyed junco. Hormones and Behavior 65, 179–187. (doi:10.1016/j.yhbeh.2013.10.006)

24. Kitano J, Kawagishi Y, Mori S, Peichel CL, Makino T, Kawata M, Kusakabe M. 2011 Divergence in Sex Steroid Hormone Signaling between Sympatric Species of Japanese Threespine Stickleback. PLoS ONE 6, e29253. (doi:10.1371/journal.pone.0029253)

25. Borg B. 2007 Reproductive Physiology of Sticklebacks. In Biology of the three-spined stickleback (eds S Östlund-Nilsson, I Mayer, F Huntingford), pp. 225–248. Boca Raton: CRC Press.

26. Crews D. 1984 Gamete production, sex hormone secretion, and mating behavior uncoupled. Hormones and Behavior 18, 22–28. (doi:10.1016/0018-506X(84)90047-3)

27. Borg B. 1982 Seasonal effects of photoperiod and temperature on spermatogenesis and male secondary sexual characters in the three-spined stickleback, *Gasterosteus aculeatus* L. Can. J. Zool. 60, 3377–3386. (doi:10.1139/z82-427)

28. Sokołowska E, Kulczykowska E. 2006 Annual reproductive cycle in two free living populations of three-spined stickleback (Gasterosteus aculeatus L.): patterns of ovarian and testicular development. Oceanologia 48, 103–121.

29. Zbinden M, Largiadèr CR, Bakker TCM. 2001 Sperm allocation in the three-spined stickleback. Journal of Fish Biology 59, 1287–1297. (doi:10.1111/j.1095-8649.2001.tb00192.x)

30. Baker JA. 1994 Life history variation in female threespne stickleback. In The evolutionary biology of the threespine stickleback (eds MA Bell, SA Foster), pp. 144–187. New York: Oxford University Press. (10.1093/oso/9780198577287.003.0006)

31. Borg B. 1994 Androgens in teleost fishes. *Comparative Biochemistry and Physiology Part C: Pharmacology*, Toxicology and Endocrinology 109, 219–245. (doi:10.1016/0742-8413(94)00063-G)

32. Hoar WS. 1962 Hormones and the reproductive behaviour of the male three-spined stickleback (Gasterosteus aculeatus). Animal Behaviour 10, 247–266. (doi:10.1016/0003-3472(62)90049-0)

33. Andersson E, Mayer I, Borg B. 1988 Inhibitory effect of 11-ketoandrostenedione and androstenedione on spermatogenesis in the three-spined stickleback, *Gasterosteus aculeatus* L. Journal of Fish Biology 33, 835–840. (doi:10.1111/j.1095-8649.1988.tb05530.x)

34. Borg B. 1981 Effects of methyltestosterone on spermatogenesis and secondary sexual characters in the three-spined stickleback (Gasterosteus aculeatus L.). General and Comparative Endocrinology 44, 177–180. (doi:10.1016/0016-6480(81)90245-8)

35. Barbasch TA, Farrar V, Behrens C, Dan U, Bell AM. 2026 Parental care in stickleback fish. Current Opinoin in Neurobiology

36. Maciejewski MF, Bell AM. 2026 An evolutionary shift to prioritizing mating over care is associated with consistently high androgen levels in male threespine stickleback. Hormones and Behavior 177, 105866. (doi:10.1016/j.yhbeh.2025.105866)

37. Blouw DM. 1996 Evolution of offspring desertion in a stickleback fish. Écoscience 3, 18–24. (doi:10.1080/11956860.1996.11682310)

38. Behrens C, Maciejewski MF, Arredondo E, Dalziel AC, Weir LK, Bell AM. 2024 Divergence in reproductive behaviors is associated with the evolutionary loss of parental care. The American Naturalist, 729465. (doi:10.1086/729465)

39. Ishikawa M, Mori S. 2000 Mating Success and Male Courtship Behaviors in Three Populations of the Threespine Stickleback. Behaviour 137, 1065–1080.

40. Bell AM, Peeke HVS. 2012 Individual variation in habituation: behaviour over time toward different stimuli in threespine sticklebacks (Gasterosteus aculeatus). Behav 149, 1339–1365. (doi:10.1163/1568539X-00003019)

41. Van Iersel JJA. 1953 An Analysis of the Parental Behaviour of the Male Three-Spined Stickleback (Gasterosteus Aculeatus L.). Behaviour 3, 1–159.

42. Sumarli A, Behrens C, Lucas G, Olufemi MJ, Bentzen P, Bell AM, Chain FJJ, Samuk K. 2025 Strong but diffuse genetic divergence underlies differentiation in an incipient species of marine stickleback. (doi:10.1101/2025.09.19.677379)

43. Blouw DM, Hagen DW. 1990 Breeding ecology and evidence of reproductive isolation of a widespread stickleback fish (Gasterosteidae) in Nova Scotia, Canada. Biological Journal of the Linnean Society 39, 195–217. (doi:10.1111/j.1095-8312.1990.tb00512.x)

44. Behrens C, Tucker ME, Julkowski K, Bell AM. 2025 Discrete genetic modules underlie divergent reproductive strategies in three-spined stickleback. Journal of Heredity, esaf086. (doi:10.1093/jhered/esaf086)

45. Samuk KM. 2016 The evolutionary genomics of adaptation and speciation in the threespine stickleback. University of British Columbia, Vancouver, BC, Canada. (doi:10.14288/1.0307145)

46. Borg B, Schoonen WGEJ, Lambert JGD. 1989 Steroid metabolism in the testes of the breeding and nonbreeding three-spined stickleback, Gasterosteus aculeatus. General and Comparative Endocrinology 73, 40–45. (doi:10.1016/0016-6480(89)90053-1)

47. Andersson E, Borg B, Goos HJTh. 1992 Temperature, but not photoperiod, influences gonadotropin-releasing hormone binding in the pituitary of the three-spined stickleback, gasterosteus aculeatus. General and Comparative Endocrinology 88, 111–116. (doi:10.1016/0016-6480(92)90199-T)

48. Chen Y, Chen L, Lun ATL, Baldoni PL, Smyth GK. 2025 edgeR v4: powerful differential analysis of sequencing data with expanded functionality and improved support for small counts and larger datasets. Nucleic Acids Research 53, gkaf018. (doi:10.1093/nar/gkaf018)

49. Larsson J. 2024 _eulerr: Area-Proportional Euler and Venn Diagrams with Ellipses_. R package version 7.0.2,.

50. Conway JR, Lex A, Gehlenborg N. 2017 UpSetR: an R package for the visualization of intersecting sets and their properties. Bioinformatics 33, 2938–2940. (doi:10.1093/bioinformatics/btx364)

51. Alexa A, Rahnenfuhrer J. 2017 topGO. (doi:10.18129/B9.BIOC.TOPGO)

52. Sayols S. 2023 rrvgo: a Bioconductor package for interpreting lists of Gene Ontology terms. microPublication Biology (doi:10.17912/micropub.biology.000811)

53. Munley KM, Hoadley AP, Alward BA. 2024 A phylogenetics-based nomenclature system for steroid receptors in teleost fishes. General and Comparative Endocrinology 347, 114436. (doi:10.1016/j.ygcen.2023.114436)

54. Huynh-Thu VA, Irrthum A, Wehenkel L, Geurts P. 2010 Inferring Regulatory Networks from Expression Data Using Tree-Based Methods. PLoS ONE 5, e12776. (doi:10.1371/journal.pone.0012776)

55. Harrington SA, Backhaus AE, Singh A, Hassani-Pak K, Uauy C. 2020 The Wheat GENIE3 Network Provides Biologically-Relevant Information in Polyploid Wheat. G3 Genes|Genomes|Genetics 10, 3675–3686. (doi:10.1534/g3.120.401436)

56. Kitano J, Kakioka R, Ishikawa A, Toyoda A, Kusakabe M. 2020 Differences in the contributions of sex linkage and androgen regulation to sex-biased gene expression in juvenile and adult sticklebacks. J of Evolutionary Biology 33, 1129–1138. (doi:10.1111/jeb.13662)

57. Olsson P-E et al. 2005 Molecular cloning and characterization of a nuclear androgen receptor activated by 11-ketotestosterone. Reprod Biol Endocrinol 3. (doi:10.1186/1477-7827-3-37)

58. Shen W-K, Chen S-Y, Gan Z-Q, Zhang Y-Z, Yue T, Chen M-M, Xue Y, Hu H, Guo A-Y. 2023 AnimalTFDB 4.0: a comprehensive animal transcription factor database updated with variation and expression annotations. Nucleic Acids Research 51, D39–D45. (doi:10.1093/nar/gkac907)

59. Barbasch TA, Abuwa VI, Carswell B, Bell AM. 2024 Managing the tradeoff between reproduction and survival requires flexibility in behaviour and gene regulation in three-spined stickleback. Proc. R. Soc. B. 291, 20242296. (doi:10.1098/rspb.2024.2296)

60. Shaw DE, Ross WD, Lambert AV, White MA. 2025 Single cell RNA-sequencing reveals no evidence for meiotic sex chromosome inactivation in the threespine stickleback fish. PLoS Genet 21, e1011875. (doi:10.1371/journal.pgen.1011875)

61. Behrens C, Maciejewski MF, Sumarli A, Lucas G, Lagunas-Robles G, Samuk K, Bell AM. 2025 Evidence of parental care as a novel reproductive isolating mechanism. bioRxiv (10.1101/2025.04.18.649590)

62. Stein LR, Trapp RM, Bell AM. 2016 Do reproduction and parenting influence personality traits? Insights from threespine stickleback. Animal Behaviour 112, 247–254. (doi:10.1016/j.anbehav.2015.12.002)

63. Sebire M, Katsiadaki I, Scott AP. 2007 Non-invasive measurement of 11-ketotestosterone, cortisol and androstenedione in male three-spined stickleback (Gasterosteus aculeatus). General and Comparative Endocrinology 152, 30–38. (doi:10.1016/j.ygcen.2007.02.009)

64. Mayer I, Borg B, Schulz R. 1990 Conversion of 11-ketoandrostenedione to 11-ketotestosterone by blood cells of six fish species. General and Comparative Endocrinology 77, 70–74. (doi:10.1016/0016-6480(90)90207-3)

65. Ren X, Chen X, Wang Z, Wang D. 2017 Is transcription in sperm stationary or dynamic? Journal of Reproduction and Development 63, 439–443. (doi:10.1262/jrd.2016-093)

66. Samuk K, Visty H, Schluter D. 2026 Genetic Divergence in the Absence of Strong Ecological Differences Between Coexisting White and Common Atlantic Marine Stickleback. Ecology and Evolution 16, e73655. (doi:10.1002/ece3.73655)

67. Borg B, Peute J, Reschke M, Hurk R. 1987 Effects of photoperiod and temperature on testes, renal epithelium, and pituitary gonadotropic cells of the threespine stickleback, Gasterosteus aculeatus L. Canadian Journal of Zoology 65, 14–19.

68. Baggerman B. 1972 Photoperiodic responses in the stickleback and their control by a daily rhythm of photosensitivity. General and Comparative Endocrinology 3, 466–476. (doi:10.1016/0016-6480(72)90177-3)

69. Behrens C, Young S, Arredondo E, Dalziel AC, Weir LK, Bell AM. 2025 The Evolutionary Loss of Paternal Care Is Associated With Shifts in Female Life-History Traits. Ecology and Evolution 15, e70497. (doi:10.1002/ece3.70497)

