## Supplemental Materials for "The gonad as a mediator of life history tradeoffs: Antagonistic hormonal pleiotropy facilitates evolutionary divergence in reproductive strategies"

### Supplemental Methods

*Validating GENIE3 GRN predictions of androgen receptor target genes*

To validate whether predicted targets of androgen receptor genes from GENIE3 gene regulatory networks are indeed responsive to androgen treatment, as would be expected if they are target genes of androgen receptor binding, we built gene regulatory networks using RNASeq count data from brains of male and female three-spined stickleback treated with 11-ketoandrostenedione (11-KA) or controls (data from Kitano et al., 2020). As 11-KA is the precursor to 11-ketotestosterone (11-KT), the primary androgen in teleosts (Borg, 1994) differential gene expression in response to 11-KA should be in part mediated by increased activation of androgen receptors, specifically the 11-KT responsive *ar2* in stickleback (Olsson et al., 2005). To test if GENIE3 accurately predicts this, we built a GRN in GENIE 3 v.1.33 (Huynh-Thu et al., 2010) with all of the transcription factors for stickleback from the Animal Transcription Factor Database 4.0 (Shen et al., 2023) defined as regulators and the brain RNASeq count data from Kitano et al. (2020). This yielded a network with >25 million links. Then, following the procedures outlined in Harrington et al. (2020), we selected the regulatory links across all transcription factors with the top 1 million highest weights. For each of the 1248 transcription factors detected in the brain RNASeq data, we then tested for hypergeometric overlap between the predicted target genes (within the top 1 million links) and the genes differentially expressed with 11-KA treatment (differential gene expression analysis as described in Kitano et al. (2020)) with an FDR correction for multiple comparisons. We found that the proportion of predicted *ar2* target genes that were also 11-KA biased genes (37/866) was significantly higher than the predicted median proportion for all transcription factors tested (one-sided Wilcoxon sign test, *p* < 0.001; Figure S1A), and *ar2* ranked 23^rd^ for lowest adjusted p-value across all 1248 hypergeometric overlap tests (p.adj < 0.001, Figure S1B). However, the predicted target genes for *ar1* were not significantly enriched for 11-KA biased genes (p.adj = 0.99), nor was the proportion of overlap with 11-KA biased genes significantly higher than the median (10/778, one-sided Wilcoxon sign test, *p* = 1). Given this, we used the top 1 million predicted links from the GENIE3 GRN to predict target genes for *ar2,* but not *ar1*, in our dataset.

**References for Supplemental Methods**

Borg, B. (1994). Androgens in teleost fishes. *Comparative Biochemistry and Physiology Part C: Pharmacology, Toxicology and Endocrinology*, *109*(3), 219–245. https://doi.org/10.1016/0742-8413(94)00063-G

Harrington, S. A., Backhaus, A. E., Singh, A., Hassani-Pak, K., & Uauy, C. (2020). The Wheat GENIE3 Network Provides Biologically-Relevant Information in Polyploid Wheat. *G3 Genes|Genomes|Genetics*, *10*(10), 3675–3686. https://doi.org/10.1534/g3.120.401436

Huynh-Thu, V. A., Irrthum, A., Wehenkel, L., & Geurts, P. (2010). Inferring Regulatory Networks from Expression Data Using Tree-Based Methods. *PLoS ONE*, *5*(9), e12776. https://doi.org/10.1371/journal.pone.0012776

Kitano, J., Kakioka, R., Ishikawa, A., Toyoda, A., & Kusakabe, M. (2020). Differences in the contributions of sex linkage and androgen regulation to sex‐biased gene expression in juvenile and adult sticklebacks. *Journal of Evolutionary Biology*, *33*(8), 1129–1138. https://doi.org/10.1111/jeb.13662

Olsson, P.-E., Berg, A. H., Von Hofsten, J., Grahn, B., Hellqvist, A., Larsson, A., Karlsson, J., Modig, C., Borg, B., & Thomas, P. (2005). Molecular cloning and characterization of a nuclear androgen receptor activated by 11-ketotestosterone. *Reproductive Biology and Endocrinology*, *3*(1). https://doi.org/10.1186/1477-7827-3-37

Shen, W.-K., Chen, S.-Y., Gan, Z.-Q., Zhang, Y.-Z., Yue, T., Chen, M.-M., Xue, Y., Hu, H., & Guo, A.-Y. (2023). AnimalTFDB 4.0: A comprehensive animal transcription factor database updated with variation and expression annotations. *Nucleic Acids Research*, *51*(D1), D39–D45. https://doi.org/10.1093/nar/gkac907

### FIGURES

#### Supplemental Figure 1.

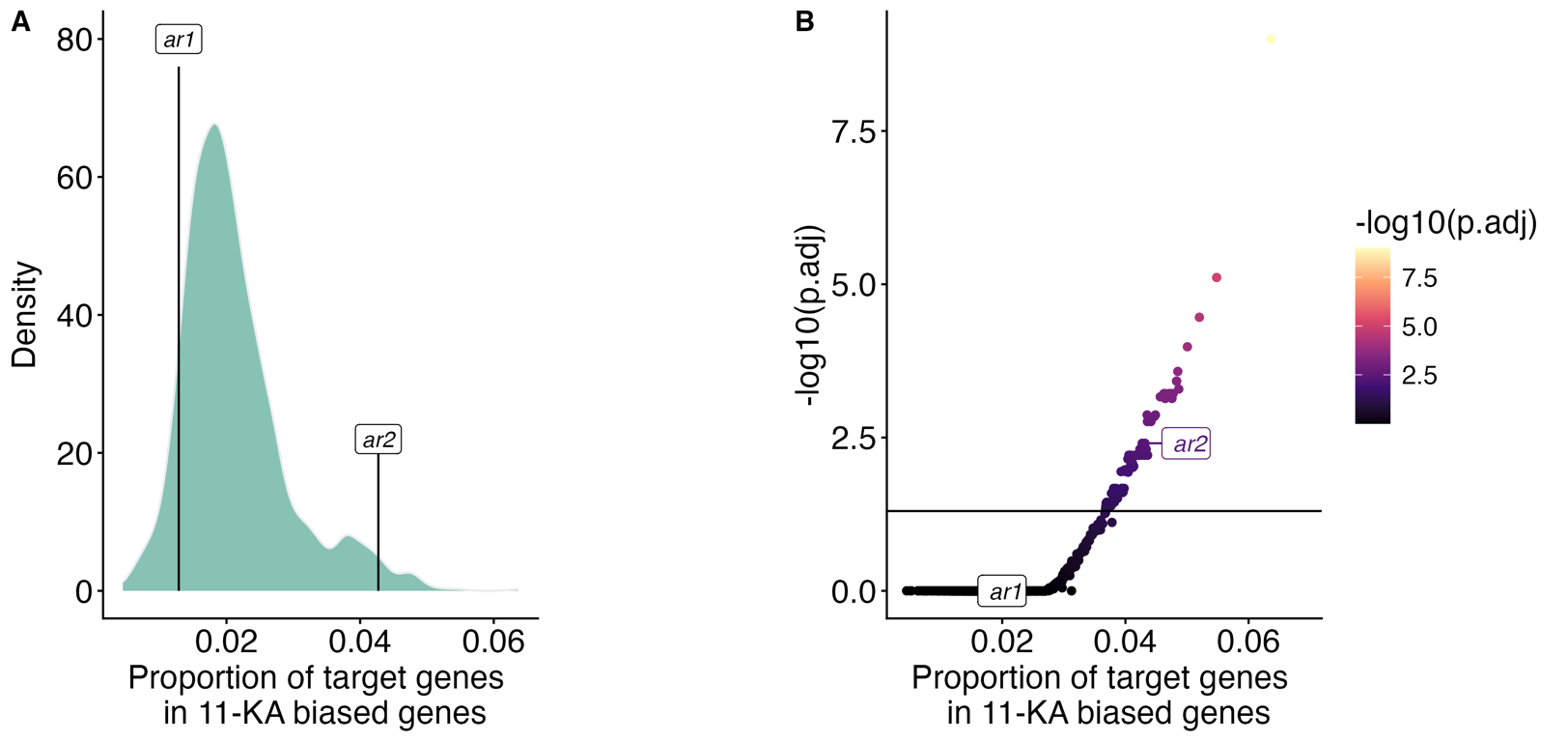

**Supplemental Figure 1.** **GENIE3 predicted targets of androgen receptor beta (*ar2*), but not androgen receptor alpha (*ar1*), are significantly enriched in genes responsive to 11-ketoandrostenedione treatment in stickleback brains** (data from (Kitano et al., 2020)). (**A**) The proportion of *ar2* target genes overlapping with 11-KA biased DEGs is significantly higher than the median overlap across the 1248 transcription factors whose predicted target genes were tested (Wilcoxon sign test, *p* < 0.01), including *ar1*. (**B**) *ar2* target genes were significantly enriched for 11-KA biased DEGs (FDR-corrected hypergeometric tests, p.adj < 0.01) and had the 23^rd^ highest ranked proportion of genes overlapping out of 1248 transcription factors tested.

#### Supplemental Figure 2.

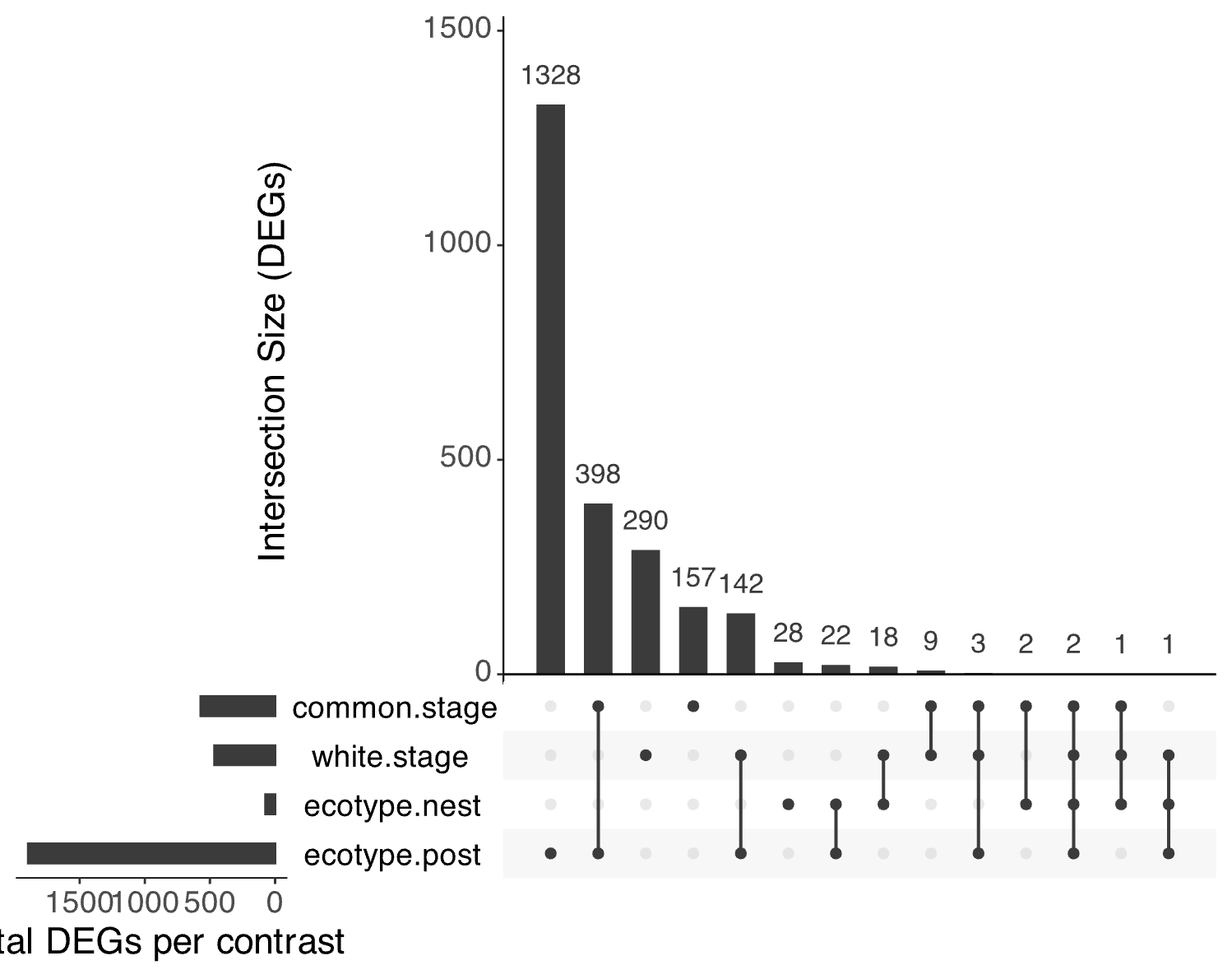

**Supplemental Figure 2.** UpSet plot showing set intersections of all pairwise differentially expressed gene (DEG) sets across pairwise contrasts. Bars to the left of the plot indicate the total number of DEGs for each contrast. Each bar indicates the number of genes shared, or unique, to the contrasts indicated with dots under each bar.

### TABLES

#### Table S1.

Candidate genes *a priori* selected for analysis based on roles in steroidogenesis and androgen regulation.

| **Gene** | **Ensembl ID** | **Description** | **Rationale for candidate gene** |
| --- | --- | --- | --- |
| *lhcgr* | ENSGACG00000034257 | luteinizing hormone chorionic gonadotropin receptor | Binding of LH to its receptor on testes activates steroidogenesis; increased sensitivity to gonadotropins may suggest increased responsiveness to HPG signals |
| *fshr* | ENSGACG00000002728 | follicle stimulating hormone receptor | Gonadotropin receptor that may stimulate gonadal steroiodgenesis or spermatogenesis |
| *star* | ENSGACG00000011782 | steroid acute regulatory protein | Activation of StAR initiates steriodogenesis via cholesterol transport; considered a rate-limiting step in steroidogenesis |
| *star2* | ENSGACG00000002322 | steroid acute regulatory protein 2 | Teleost paralog of StAR |
| *cyp11a2* | ENSGACG00000004713 | cytochrome p450 family 11 subfamily a | Converts cholesterol to pregnenolone; considered a rate-limiting step in steroiodogenesis |
| *cyp17a1* | ENSGACG00000002550 | cytochrome p450 family 17 subfamily a | Converts pregnenolone to DHEA |
| *hsd3b1* | ENSGACG00000001425 | 11-beta-hydroxysteroid dehydrogenase family 3 | Converts DHEA to androstenedione (A4) |
| *cyp11c1* | ENSGACG00000011657 | cytochrome p450 family 11 subfamily c | Converts androstenedione to 11-beta-hydroxy-androstenedione (11β-OH-A4); teleost homolog of mammalian CYP11B1 (Nelson et al. 2013) |
| *cyp19a1a* | ENSGACG00000016742 | cytochrome P450, family 19, subfamily A, polypeptide 1a | Aromatase; converts testosterone to 17β-estradiol (note: not detected in testes RNASeq data) |
| *srd5a1* | ENSGACG00000008961 | steroid-5-alpha-reductase, alpha polypeptide 1 | Converts testosterone to dihydrotestosterone (note: not detected in testes RNASeq data) |
| *hsd11b2* | ENSGACG00000017514 | hydroxysteroid 11-beta dehydrogenase 2 | Converts 11β-OH-A4 to 11-ketoandrostenedione (11-KA) |
| *hsd17b3* | ENSGACG00000006978 | hydroxysteroid 17-beta dehydrogenase 3 | Converts 11-KA to 11-KT (evidence from zebrafish) |
| *ar1* | ENSGACG00000018525 | androgen receptor 1 (alpha) | Receptors for circulating androgens |
| *ar2* | ENSGACG00000020332 | androgen receptor 2 (beta) | Receptors for circulating androgens; ar2 preferentially binds 11-ketotestosterone in stickleback (Olsson et al. 2005) |

#### Table S2.

Bonferroni-corrected hypergeometric tests between predicted targets of androgen receptor 2 and DEGs for each pairwise contrast.

| **Contrast** | **Regulator** | **DEGs in contrast (n)** | **Targets genes of regulator (n)** | **Overlap (n)** | **p** | **p.adj** |
| --- | --- | --- | --- | --- | --- | --- |
| Common-biased genes at prolonged-summer | *ar2* | 1391 | 1368 | 179 | <0.001 | <0.001 |
| White-biased genes at prolonged summer | *ar2* | 505 | 1368 | 54 | <0.001 | 0.001 |
| Commons: breeding vs. prolonged summer | *ar2* | 572 | 1368 | 29 | 0.934 | 1 |
| Whites: breeding vs. prolonged summer | *ar2* | 466 | 1368 | 42 | 0.019 | 0.114 |
| Whites vs. commons: breeding | *ar2* | 74 | 1368 | 2 | 0.957 | 1 |
| Whites vs. commons: prolonged summer | *ar2* | 1896 | 1368 | 233 | <0.001 | <0.001 |

#### Table S3.

Gene ontology (GO) terms significantly enriched (Fisher’s exact test, p < 0.01) in 1896 DEGs between ecotypes under prolonged summer conditions. Cluster and parent term are assigned to reduce GO terms by semantic similarity using *rrvgo*.

#### Table S4

Gene ontology (GO) terms significantly enriched (Fisher’s exact test, p < 0.01) in 74 DEGs between ecotypes during breeding. Cluster and parent term are assigned to reduce GO terms by semantic similarity using *rrvgo*.

**Table S5.**

Gene ontology (GO) terms significantly enriched (Fisher’s exact test, p < 0.01) in genes upregulated, or biased, towards commons under prolonged summer conditions. Cluster and parent term are assigned to reduce GO terms by semantic similarity using *rrvgo*.

**Table S6.**

Gene ontology (GO) terms significantly enriched (Fisher’s exact test, p < 0.01) in genes upregulated, or biased, towards whites under prolonged summer conditions. Cluster and parent term are assigned to reduce GO terms by semantic similarity using *rrvgo*.
